# Replication of seizure-suppressing effects of alpha-linolenic acid on the *Drosophila melanogaster para*^*Shudderer*^ mutant

**DOI:** 10.1101/2025.08.20.671359

**Authors:** Samantha Dinnel, Brady Weimer, Allyson Davis, Katie Gerovac, Leah M Hawbaker, Matthew Maser, Meghan Shannon, Douglas J Brusich

**Author notes:** <u>**Corresponding Author:**</u> Dr. Douglas J Brusich. authors contributed equally to this work (Samantha Dinnel, Brady Weimer). <u>**Author Contributions:**</u> **Samantha Dinnel**: Investigation, Validation, Writing – Review and Editing **Brady Weimer**: Investigation, Validation, Writing – Review and Editing **Allyson Davis**: Investigation, Validation, Writing – Review and Editing **Katie Gerovac**: Investigation, Validation, Writing – Review and Editing **Leah Hawbaker**: Investigation, Validation, Writing – Review and Editing **Matthew Maser**: Investigation, Validation, Writing – Review and Editing **Meghan Shannon**: Investigation, Validation, Writing – Review and Editing **Douglas J Brusich**: Conceptualization, Data Curation, Formal Analysis, Funding Acquisition, Investigation, Methodology, Project Administration, Resources, Supervision, Validation, Visualization, Writing – Original Draft, Writing – Review and Editing. <u>**Funding:**</u> UWLAX Start-up.

## Abstract

Voltage-gated sodium channels are essential for healthy nervous system function. Mutations in voltage-gated sodium channels are associated with a range of seizure conditions. The genetics of seizure conditions are complex and often challenging to study or replicate in animal models. The *Drosophila melanogaster* gene *paralytic* (*para*) is the sole voltage-gated sodium channel gene in flies. The *para*^*Shudderer*^ allele causes dominant seizure activity manifest in adult morphology and behavior. In this study we replicated previous findings of *para*^*Shudderer*^ hyperexcitability and the ability to suppress this activity with dietary supplementation of an omega-3 fatty acid, alpha-linolenic acid (ALA). Our results support the robustness of the *para*^*Shudderer*^ phenotype and the replicability of findings across separate lab environments.

## Description

The voltage-gated sodium channel Na_V_1.1 is central to the function of inhibitory neurons of the mammalian central nervous system (CNS) (Ogiwara et al. 2007; Ricobaraza et al. 2023). Accordingly, mutations in the gene encoding the pore-forming subunit of the Na_V_1.1 complex (*SCN1A*) and resultant channel dysfunction are associated with a range of epileptic and neurodevelopmental conditions including Dravet Syndrome, generalized epilepsy with febrile seizures plus (GEFS+), familial hemiplegic migraine type-2, and developmental and epileptic encephalopathy (Catterall et al. 2010; Meisler et al. 2010). The genetics of these disorders are complicated by the number of *SCN1A* mutations associated with disease, such as the >500 *SCN1A* mutations associated with Dravet Syndrome, and incomplete penetrance of disease-associated alleles (Bonanni et al. 2004; Mantegazza et al. 2005; Marini et al. 2011).

The fruit fly (*Drosophila melanogaster*) has long-served as an experimental model for epilepsy . Fruit flies have a single voltage-gated sodium channel ortholog (*paralytic*) and flies have been used to model Dravet Syndrome and GEFS+ (Suzuki et al. 1971; Sun et al. 2012; Schutte et al. 2014; Tapia et al. 2021). The *Shudderer* mutation was first discovered in 1971 for its spontaneous seizure activity and was later found to be a dominant allele of *paralytic* (*para*^*Shudderer*^ hereafter *para*^*Shu*^) resulting from a methionine to isoleucine amino acid substitution in the first transmembrane segment of the third domain of the protein (Williamson 1982; Kaas et al. 2016). *para*^*Shu*^ causes spontaneous seizure activity, a wings-apart morphology consistent with hyperexcitability, and high temperature seizure activity (Kaas et al. 2016). The hyperexcitable seizure behavior of *para*^*Shu*^ can be suppressed by dietary supplementation with milk whey or omega-3 fatty acids more specifically, genetic downregulation of glutathione S-transferase S1 (*GstS1*), or removal of the resident gut microbiota (Kasuya et al. 2019; Chen et al. 2020; Kasuya et al. 2023; Lansdon et al. 2024).

We first aimed to replicate the core findings of *para*^*Shu*^ hyperexcitability. We used a *para*^*Shu*^*/Fm7* balancer stock from previous work or the same *para*^*Shu*^ allele rebalanced over *Fm7A-Tb* (Kaas et al. 2016). Each line was crossed to *Canton-S* wild-type male flies to generate *para*^*Shu*^/+ experimental progeny. We found no discernable difference in the rates of wings-apart morphology nor adult temperature-sensitive (TS) seizure activity amongst these *para*^*Shu*^/+ progeny (wings-apart: p = 0.63 by Fisher’s Exact test of n ≥ 123 flies per dataset; TS activity: p = 0.24 for genotype differences by Two-Way repeated measures ANOVA with n ≥ 3 datasets of at least 30 flies each).

We next sought to replicate the suppressive effects of dietary alpha-linolenic acid (ALA) supplementation on *para*^*Shu*^/+ hyperexcitable phenotypes. Data from crosses of *para*^*Shu*^*/Fm7* and *para*^*Shu*^*/Fm7A-Tb* to *Canton-S* were combined into a merged dataset for these investigations. Blue dye was added to standard food along with ALA to ensure uniformity of mixing, and we therefore also included a standard diet with blue dye only control condition. Wild-type *Canton-S* flies showed no evidence of either wings-apart morphology or TS seizure activity (Figs. 1A/B) across each of the standard diet (+), blue dye control diet (+Blue), or the standard diet supplemented with blue dye and 0.05% (1.8 mM) ALA (+Blue, +ALA). By contrast, *para*^*Shu*^/+ flies on standard or blue dye control food each exhibited similarly significant levels of aberrant wing morphology (∼80% wings apart) compared to *Canton-S* wild-type controls (Fig. 1A). Consistent with previous findings, we found *para*^*Shu*^/+ animals fed a diet supplemented with 0.05% ALA had significantly reduced levels of aberrant wing morphology compared to the dietary controls (Fig. 1A) (Kasuya et al. 2023).

**Figure 1:**
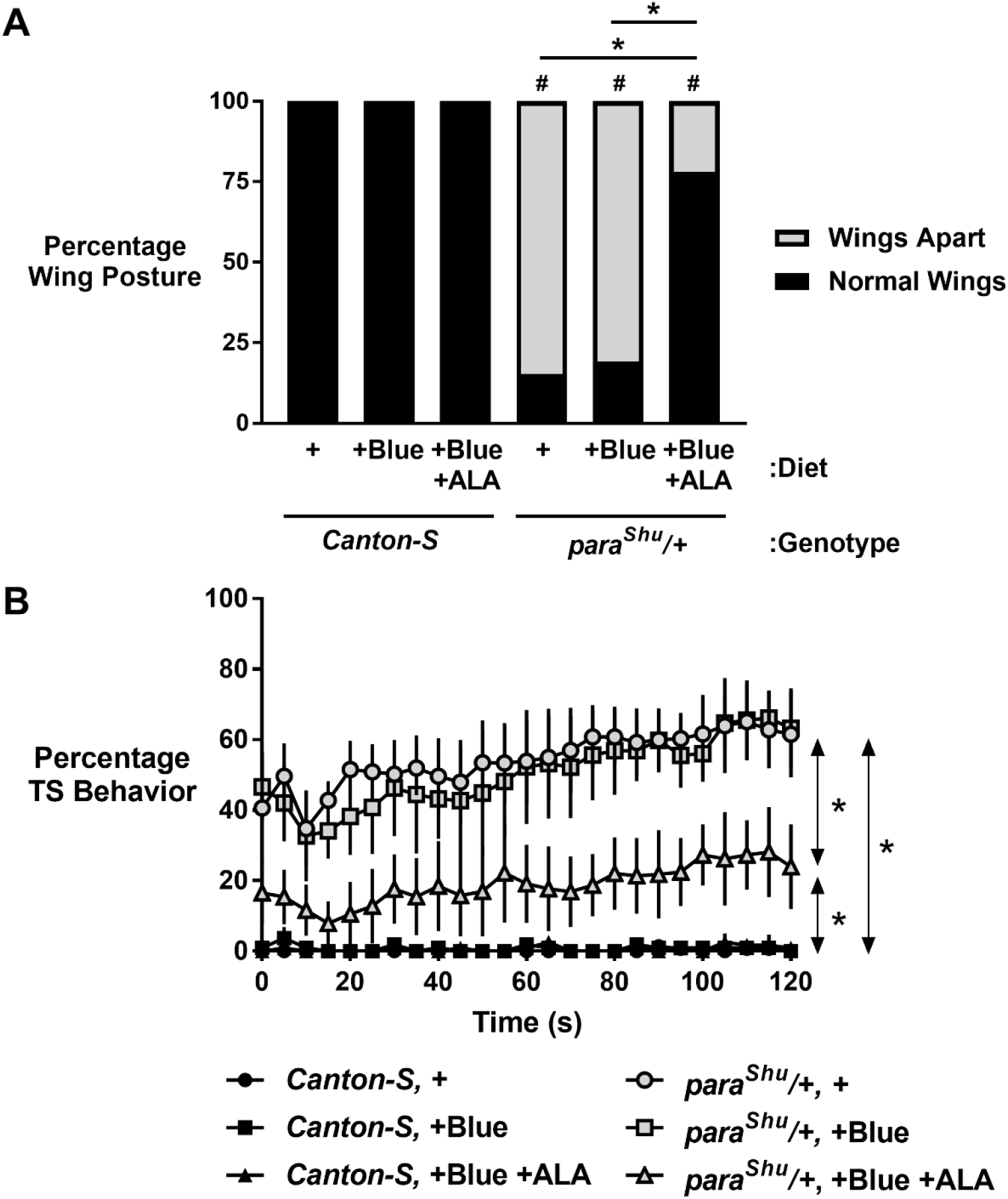
ALA suppresses *para*^*Shu*^/+ adult hyperexcitability. (A) *para*^*Shu*^/+ adult wings-apart morphology is suppressed by 0.05% ALA. # signifies a difference compared to the respective *Canton-S* control and * signifies a difference between indicated datasets by Fisher’s Exact Test with Bonferroni correction (p < 0.0001). Diets are standard (+), standard with blue dye (+Blue), or standard with blue dye and 0.05%ALA (+Blue, +ALA). n ≥ 180 flies per condition. (B) *para*^*Shu*^/+ adult temperature-sensitive (TS) seizure behavior is suppressed by 0.05% ALA. * = p < 0.05 between conditions by Two-Way ANOVA with post-hoc comparison between conditions. Diet conditions as in (A). n ≥ 3 datasets of at least 30 flies each. Data plotted are means with standard deviations.

We similarly found that ALA suppressed adult TS seizure activity. Wild-type flies showed little to no evidence of locomotor dysfunction across 2 minutes at 37°C regardless of diet (Fig. 1B). However, *para*^*Shu*^/+ flies showed pronounced seizure activity when reared on food lacking ALA, and this phenotype was significantly, but not entirely, ameliorated when *para*^*Shu*^/+ animals were raised on 0.05% ALA-containing food (Fig. 1B).

Our results replicate previous findings of seizure activity in *para*^*Shu*^/+ adults, and the ability of dietary supplementation of an omega-3 fatty acid, alpha-linolenic acid (ALA), to suppress this hyperexcitability (Williamson 1982; Kaas et al. 2016; Kasuya et al. 2023). These findings confirm the robustness of the *para*^*Shu*^/+ phenotype across separate genetic backgrounds (*para*^*Shu*^/+ in a *Fm7* or *Fm7-Tb* background) and lab environments. Dietary lipids or ALA must be present during developmental larval stages for suppression of adult hyperexcitability (Kasuya et al. 2019; Kasuya et al. 2023). Future studies will investigate if ALA is capable of suppressing larval hyperexcitability, or, as with acute lithium administration, it is ineffective at this stage (Kaas et al. 2016). Overall, our findings support the utility and replicability of the *Drosophila* system and *para*^*Shu*^/+ allele for studies of seizure disorders.

## Methods

### Fly Husbandry

*para*^*Shudderer*^*/Fm7* and *Canton-S* were generously provided by Atulya Iyengar (University of Alabama). *para*^*Shudderer*^*/Fm7* was rebalanced over *Fm7A-Tb* (BDSC 36489). *para*^*Shudderer*^*/Fm7 and para*^*Shudderer*^*/Fm7A-Tb* were maintained at 23°C on a 12H:12H light:dark cycle. Virgin females from *para*^*Shudderer*^ stocks were crossed to *Canton-S* males to generate *para*^*Shudderer*^/+ experimental females to match prior studies examining phenotypes and suppression with alpha-linolenic acid from prior studies (Kaas et al. 2016; Kasuya et al. 2019; Kasuya et al. 2023) *Canton-S* was handled similarly to generate *Canton-S* control females. All experimental crosses were maintained at 25°C on the same 12H:12H light:dark cycle. Flies were maintained on a standard glucose, agar, cornmeal, yeast diet with the following compositions per liter of deionized water: 7.66 g agar (Genesee), 14.4 g D-glucose (Dot Scientific), 50.3 g cornmeal (Genesee), 15 g inactive dry yeast (Genesee), 5.66 ml 10% tegosept (Andwin Scientific), 4.68 ml propionic acid (99.5%; Thermo Scientific), and 0.46 ml phosphoric acid (85%; Fisher Chemical). Subsets of experimental animals were raised on standard food supplemented with FD&C no. 1 brilliant blue dye (Fisher Scientific) at 1 µg/ml (+Blue diet) or this blue dye diet with 0.05% (1.8 mM) alpha-linolenic acid (Nu-Chek Prep; +Blue, +ALA diet).

### Wings-apart and Adult Temperature-Sensitive Testing

Experimental animals were collected on CO_2_ within 5 days of eclosion and scored for wing posture (normal or wings-apart) as reported previously (Kaas et al. 2016). Following collection, animals were allowed a minimum of 24 hours post-CO_2_ to recover. Animals aged 1-5 days after eclosion were tested for temperature-sensitive (TS) behavior after dividing animals to a density of 1-2 animals per glass vial. Vials were immersed in a 37°C water bath for 2 minutes and scored every 5 seconds for TS behavior by scoring loss of normal standing posture, which included paralysis and convulsion events (Kaas et al. 2016).

### Statistics

All statistical testing and data visualization was performed using GraphPad Prism v7.05 (GraphPad Software, La Jolla California USA). Wing posture was assessed by Fisher’s exact test of count data with Bonferroni correction for multiple testing (α = 0.005). Temperature-sensitive seizure data were arrayed with condition (genotype + diet) as columns and time-points as rows. Each dataset was composed of at least 3 independent replicates of 30 or more flies. Temperature-sensitive seizure data were statistically tested by two-way ANOVA with multiple comparisons for a mean column effect and Holm-Sidak correction for multiple comparisons (α = 0.05).

